# Glucosinolate catabolism maintains glucosinolate profiles and transport in sulfur-starved *Arabidopsis*

**DOI:** 10.1101/2022.12.21.521326

**Authors:** Liu Zhang, Ryota Kawaguchi, Takuro Enomoto, Sho Nishida, Meike Burow, Akiko Maruyama-Nakashita

## Abstract

Glucosinolates (GSL) are sulfur (S)-rich specialized metabolites produced by plants of the Brassicales order. Our previous study found that in Arabidopsis seedlings, S deficiency (−S) promoted GSL catabolism by activating two ß-glucosidases (BGLU), BGLU28 and BGLU30. The induced GSL catabolism was a survival strategy for seedlings grown under −S, because S released from GSL was reincorporated into primary S metabolites which are essential for plant growth. However, as GSL profile in plants vary among growth stages and organs, we set out to test a potential contribution of BGLU28/30-dependent GSL catabolism at the reproductive growth stage. Thus, in this study, we assessed growth, metabolic, and transcriptional phenotypes of mature *bglu28/30* double mutants grown under different S conditions. Our results showed that compared to wild-type plants grown under −S, mature *bglu28/30* mutants displayed impaired growth and accumulated increased levels of GSL in their mature seeds, siliques, flowers, and rosette leaves of before bolting plants. In contrast, the levels of primary S-containing metabolites, glutathione and cysteine, were decreased in mature seeds. Furthermore, the transport of GSL from rosette leaves to the reproductive organs was stimulated in the *bglu28/30* mutants under −S. Transcriptome analysis revealed that genes related to other biological processes, such as phytohormone signaling and plant response to heat, responded differentially to −S in the *bglu28/30* mutants. Altogether, these findings broadened our understanding of the roles of BGLU28/30-dependent GSL catabolism in plant adaptation to nutrient stress.

**One-sentence summary:** Disruption of glucosinolate catabolic genes, *BGLU28* and *BGLU30*, in sulfur-starved mature *Arabidopsis* impaired growth, affected glucosinolate distribution, and altered transcriptional profiles.

## Introduction

Specialized metabolites in the Brassicales order, glucosinolates (GSL), have attracted wide attention from researchers for their defensive function in plants and medicinal values for humans (Halkier and Gershenzon, 2006; Wittstock and Burow, 2012; Abdull Razis and Mohd Noor, 2013; Maina et al., 2020). Interestingly, recent studies have revealed that GSL and their breakdown products have more physiological roles than expected. These natural products are reported to be signaling molecules to mediate root growth, flowering time, and be involved in drought stress induced stomatal aperture regulation (Zhao et al., 2008; Kerwin et al., 2011; Jensen et al., 2015; Katz et al., 2015; Malinovsky et al., 2017; Salehin et al., 2019; Katz et al., 2020). As sulfur (S)-containing compounds, GSL makes up around 30 % of organic S in Brassica plants and were found to serve as the S storage molecules, which can be reintegrated into primary metabolites upon hydrolysis under S deficiency (−S) (Zhang et al., 2020; Sugiyama et al., 2021).

GSL are derived from amino acid precursors; the structurally diverse GSL compounds can be divided into three groups: aliphatic (mGSL), indolic (iGSL), and benzenic GSL (Grubb and Abel, 2006; Halkier and Gershenzon, 2006; Blažević et al., 2020). The bioactivities of GSL are mainly attributed to their catabolism catalyzed by a specialized group of ß-glucosidases (BGLUs) named myrosinases. Among the 47 *Arabidopsis thaliana* BGLUs, 22 have been presumed to be myrosinases and are divided into two groups, classical myrosinases (BGLU34-BGLU39) and atypical myrosinases (BGLU18-BGLU33), based on their amino acid sequence signatures (Bednarek et al., 2009; Nakano et al., 2017). The classical BGLUs, BGLU38/TGG1, and BGLU37/TGG2, are both expressed in the above-ground tissues and contribute to plant defense against insects by mainly catalyzing aliphatic GSL catabolism (Barth and Jander, 2006). BGLU38/TGG1 is the most abundant protein in guard cells and plays a role in plant stomatal defense against bacterial pathogens (Zhao et al., 2008; Zhang et al., 2019). BGLU34/TGG4 and BGLU35/TGG5 are expressed in root tips where they are involved in the regulation of root growth possibly through converting iGSL to a precursor of auxin (Fu et al., 2016; Hornbacher et al., 2022). BGLU39/TGG3 and BGLU36/TGG6 are both pseudogenes in *Arabidopsis* Col-0 but show tissue-specific expression in reproductive organs (Zhang et al., 2002; Wang et al., 2009). As for the atypical BGLUs, BGLU26/PEN2 was the first identified atypical BGLU that contributes to plant antifungal defense by favorably catalyzing iGSL hydrolysis (Bednarek et al., 2009; Clay et al., 2009). BGLU23/PYK10 and BGLU18 are accumulated in ER bodies, and they provide an effective defense against herbivore attacks by degrading a specific iGSL, 4-methoxyindole-3-yl-methylglucosinolate (4MI3G) (Matsushima et al., 2003; Ogasawara et al., 2009; Nakazaki et al., 2019).

Notably, the roles of another two atypical BGLUs, BGLU28 and BGLU30 (BGLU28 /30), have long been proposed in plant response to −S due to the up-regulation of their transcript levels and the drastic decrease of GSL levels under this nutrient stress (Maruyama-Nakashita et al., 2003, 2006; Nikiforova et al., 2003; Hirai and Saito, 2004; Maruyama-Nakashita, 2017). The reduction of mGSL levels under −S was also attributed to the repression of their biosynthesis by the action of Sulfur Deficiency Induced1 (SDI1) and SDI2. These two regulatory genes are induced by −S to repress the transactivation activity of the R2R3 MYB transcription factor, MYB28, towards mGSL biosynthetic genes (Gigolashvili et al., 2007; Hirai et al., 2007; Aarabi et al., 2016). However, disruption of *SDI1* and *SDI2* cannot fully recover mGSL levels under −S to that in the wild-type plants (WT) under S sufficiency (+S), which suggested the contribution of GSL catabolism to reduce GSLs level under −S (Aarabi et al., 2016). Latest studies on −S-induced GSL catabolism and the potential role of GSL as S storage molecules provided pivotal proof to support the hypotheses that 1) BGLU28 and BGLU30 play the main role in catalyzing GSL degradation under –S, and 2) this catabolic process enables the retrograde flow of S from GSL to primary S-containing metabolites, thus sustaining plant growth (Zhang et al., 2020; Sugiyama et al., 2021). These findings prompted us to further characterize the functions of these two –S-inducible BGLUs. As the previous studies only analyzed plant seedlings, GSL catabolism in the mature plants grown under –S still awaits investigation.

GSL composition and content are known to vary strongly between plant tissues as well as in response to –S. The reproductive tissues, seeds, and siliques show the highest GSL diversity and concentrations, whereas roots display the lowest diversity, and senescent rosette leaves contain the lowest GSL levels (Petersen et al., 2002; Brown et al., 2003). The highest concentrations of GSL are found in seeds are maintained even under −S (Morikawa-Ichinose et al., 2019). Such tissue-specific distribution of GSL is presumed to result from the coordination of GSL catabolism, biosynthesis, and long-distance transport facilitated by GSL transporters, such as GSL Transporter1 (GTR1) and GTR2 (Nour-Eldin et al., 2012; Andersen et al., 2013; Jeschke et al., 2019). However, little is known about how the distribution of GSL in adult plants changes under S stress and what the significance of these changes is for plant fitness.

In this study, we took advantage of *bglu28/30* mutants to investigate GSL catabolism in mature plants under –S. Our results suggest that the BGLU28/30-dependent GSL catabolism occurs in the reproductive organs and rosette leaves under –S. This catabolic process sustains growth and reproduction when S is limiting. Furthermore, disruption of *BGLU28*/*30* also affected GSL translocation among organs and was associated with tissue-specific transcriptional changes in plants grown under –S. Overall, our results broadened our knowledge about these two atypical BGLUs and emphasized the physiological significance of GSL catabolism for plants grown under S stress.

## Results

### Disruption of *BGLU28* and *BGLU30* impaired the growth performance of mature plants under −S

To examine whether the activity of BGLU28 and BGLU30 contributes to overall plant growth, we observed the growth of mature WT and *bglu28/30* grown in vermiculite supplied with sufficient (+S, 750 µM) and deficient (−S, 0 µM) levels of sulfate (Figure 1, A and B). Except for the decrease in the fresh weight of *bglu28/30’*s rosette leaves, these two genotypes showed similar growth status under +S (Figure 1C). But under −S, in addition to the reduced rosette leaves’ fresh weight, the aerial parts’ dry weight, primary stem height, and seed yield of *bglu28/30* mutants were all decreased compared to WT (Figure 1C). These observations indicated important roles of BGLU28 and BGLU30 in maintaining plant growth and development under S stress.

**Figure 1.**
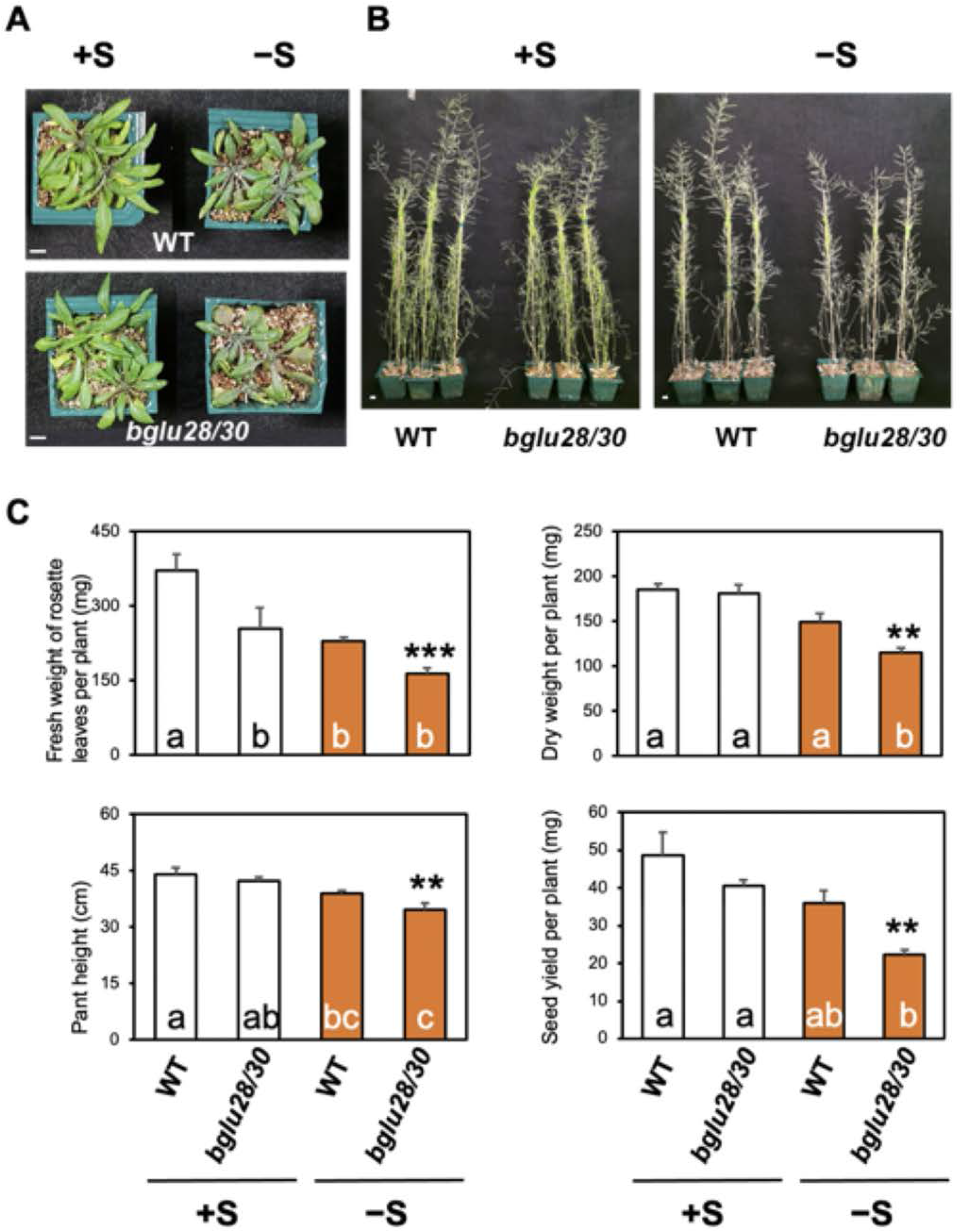
Growth phenotypes of mature *bglu28/30* and the wild-type (WT) plants under different S conditions. Plants were grown on vermiculite supplied with sufficient S (+S, white bar) or deficient S (−S, orange bar). (A) and (B) Representative images of the rosette leaves of 5-week-old plants and the whole 6-week-old plants grown on vermiculite. Scale bar, 1 cm. (C) Fresh weight of rosette leaves per 5-week-old plant (n =4). Dry weight of the aerial part per plant after seed harvesting (n = 4). Primary stem height per 6-week-old plant (n = 12). Dry weight of seeds harvested from one plant (n = 4). The values and error bars indicate means ± SEM. Two-way ANOVA followed by the Tukey–Kramer test was applied to all experimental conditions. Different letters indicate significant differences (*P* < 0.05). Asterisks indicate significant difference detected by Student’s *t*-test between WT and *bglu28/30* mutants under the same S condition (** 0.01< *P* <0.05, *** *P*<0.01).

### *BGLU28* and *BGLU30* have distinct tissue-specific expression patterns under −S

To better characterize the function of *BGLU28* and *BGLU30* under S stress, we amplified 2928 bp and 2045 bp upstream regions of *BGLU28* and *BGLU30*, respectively, and fused the fragments with the coding sequence of yellow fluorescent protein (YFP). The constructs were then introduced into WT plants and named *P*_*BGLU28*_-*YFP* and *P*_*BGLU30*_-*YFP*, respectively. *T*_*2*_ generations of these transgenic and WT plants were grown under different S conditions, and YFP fluorescence was observed (Supplemental Figure S1; Figure 2).

**Figure 2.**
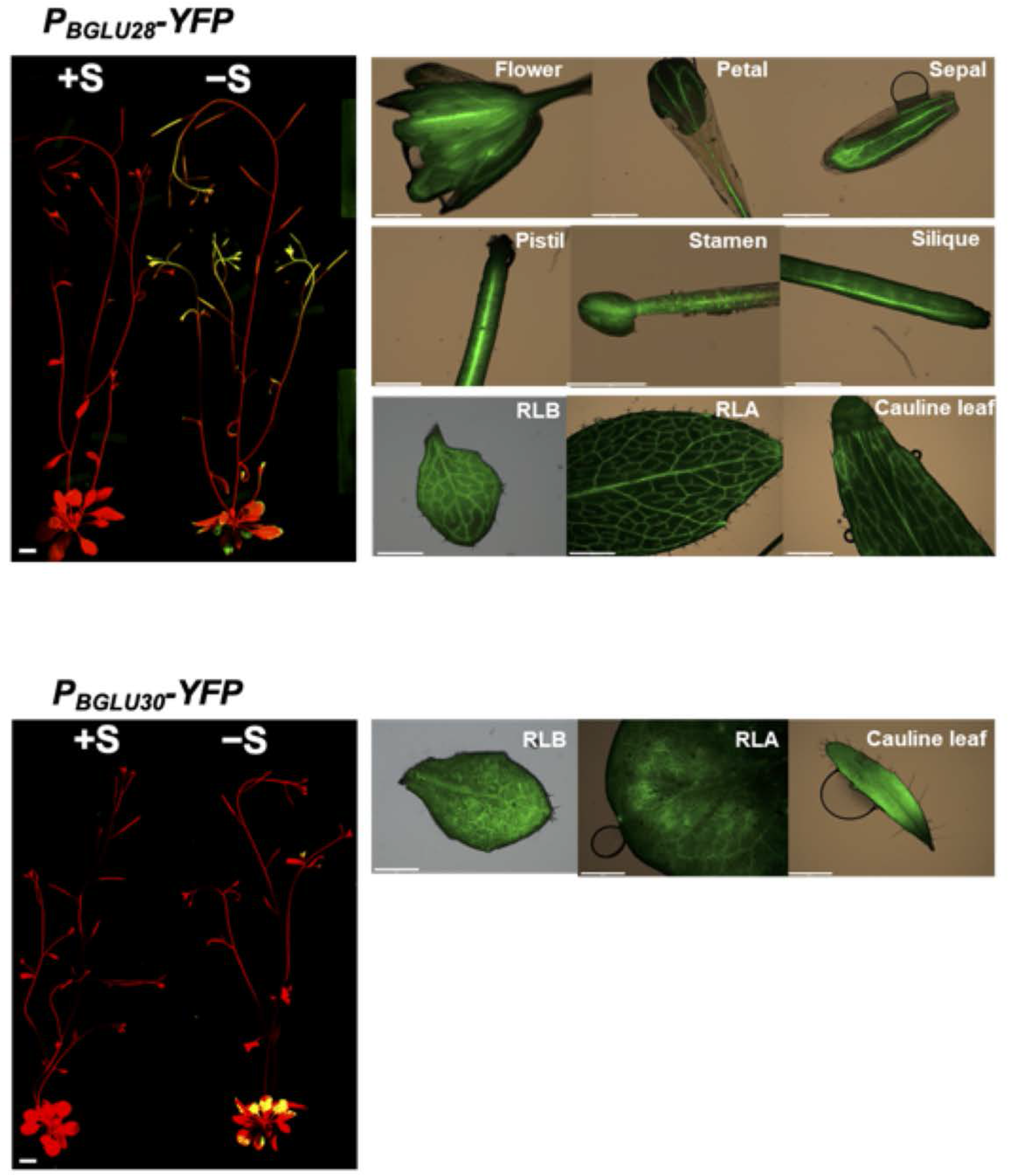
Tissue-specific expression patterns of *BGLU28* and *BGLU30* under −S. *P*_*BGLU28*_-*YFP* and *P*_*BGLU30*_-*YFP* expression pattern in the whole 5-week-old mature plants grown under both +S and −S (left panel) and in the specific organs of 4-week-old (before bolting) and 5-week-old (after bolting) plants grown under −S (right panel). Scale bar, 1 cm: the whole mature plants; 1500 µm: rosette leaf of before-bolting (RLB) and after-bolting (RLA) plants, cauline leaf; 750 µm: flower, petal, sepal, pistil, silique; 500 µm: stamen.

No fluorescence can be observed in WT plants and the YFP signals driven by *BGLU28* and *BGLU30* promoters in mature plants were only observed under −S (Supplemental Figure S1; Figure 2). The fluorescence signal from *BGLU28* promoter was extensively detected in the rosette leaves, the cauline leaves, and the signal was extremely intense in the flowers, developing siliques, and the apical region of stems (Figure 2). Unlike the extensive expression of *BGLU28* in various organs, *BGLU30* was only expressed in mature plants’ rosette leaves and young cauline leaves. Besides, unlike the vascular tissue-specific expression of *BGLU28* in the leaves, *BGLU30* was distributed in the whole leaf blades (Figure 2).

### The disruption of *BGLU28* and *BGLU30* alters glucosinolates (GSL) content in different plant tissues

The expression of *BGLU28* and *BGLU30* in diverse organs of mature plants urged us to analyze whether these two BGLUs modified GSL profile in these organs under −S. Thus, we extracted GSL from the mature seeds, developing siliques, flowers, and rosette leaves of WT and *bglu28/30* grown under different S conditions. Seven major GSL detected in *Arabidopsis* were analyzed, which include six mGSL: 3-methylsulfinylpropyl GSL (3MSOP), 4-methylsulfinylbutyl GSL (4MSOB), 8-methylsulfinyloctyl GSL (8MSOO), 4-methylthiobutyl GSL (4MTB), 7-methylthioheptyl GSL (7MTH), 8-methylthiooctyl GSL (8MTO), and one iGSL, indol-3-yl-methyl GSL (I3M) (Figure 3). These six mGSL can be classified into two groups depending on their structures: methylsulfinylalkyl (MSOX) GSL that consists of 3MSOP, 4MSOB, 8MSOO, and methylthioalkyl (MTX) GSL that are made up of 4MTB, 7MTH, 8MTO. We found that MTX GSL is much more dominant in the seeds, while MSOX GSL is more abundant in other tissues (Figure 3).

**Figure 3.**
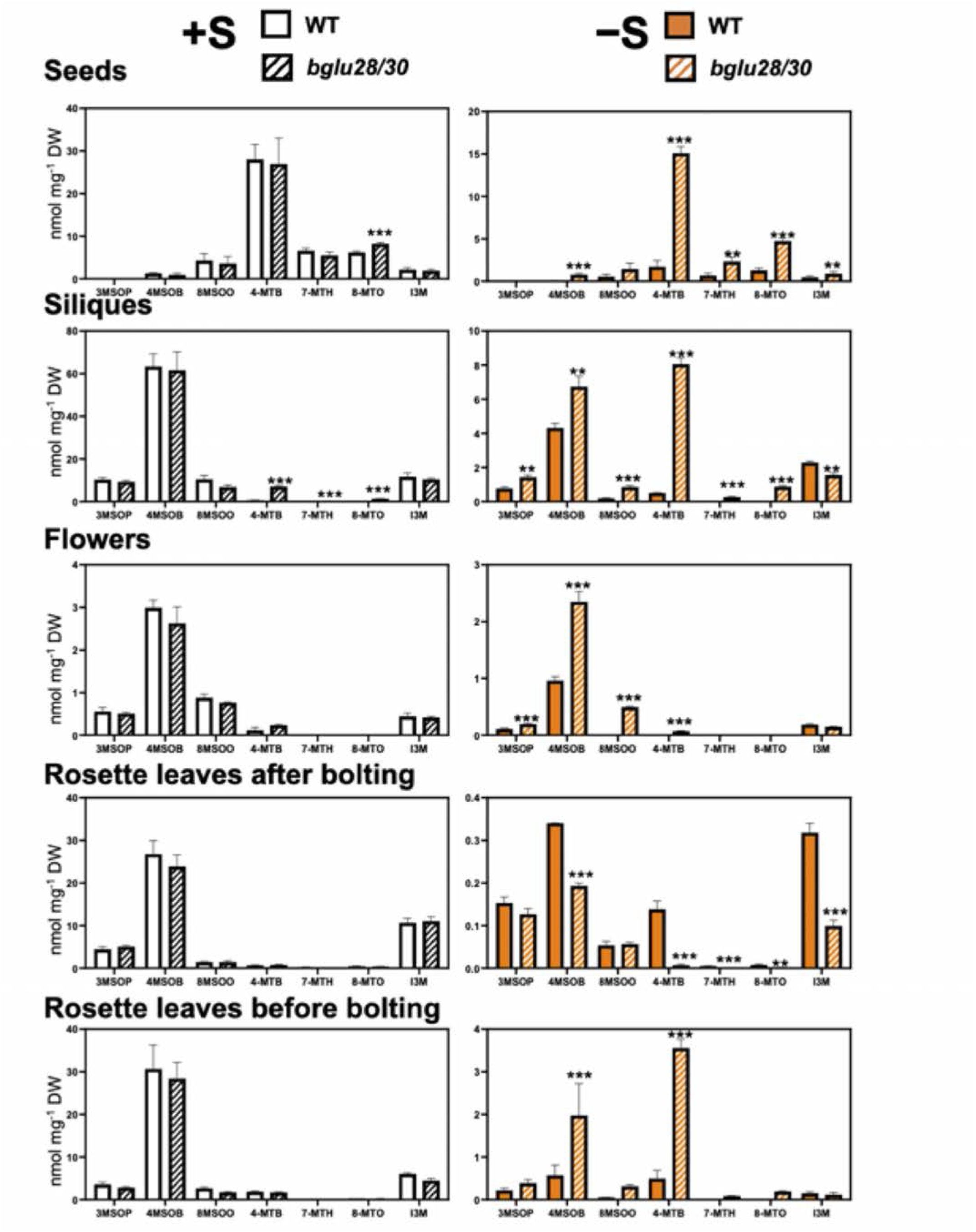
GSL content in different parts of *bglu28/30* and the wild-type (WT) plants grown under different S conditions. Plants were grown on vermiculite supplied with sufficient S (+S, white bar: WT plants, bar with black diagonal lines: *bglu28/30* mutants) or deficient S (−S, orange bar: WT plants, bar with orange diagonal lines: *bglu28/30* mutants) for 4 weeks (before bolting) and 5 weeks (after bolting). The content of 3-methylsulfinylpropyl GSL (3MSOP), 4-methylsulfinylbutyl GSL (4MSOB), 8-methylsulfinyloctyl GSL (8MSOO), 4-Methylthiobutyl GSL (4MTB), 7-methylthioheptyl GSL (7MTH), 8-methylthiooctyl GSL (8MTO), and indol-3-ylmethyl GSL (I3M) per mg dry weight (DW) of each plant tissue were analyzed by LC-MS. The values and error bars indicate means ± SEM (n = 3). Asterisks indicate significant difference detected by Student’s *t*-test between WT and *bglu28/30* under the same S condition (** 0.01< *P* <0.05, *** *P*<0.01).

Although the expression of *BGLU28* and *BGLU30* was not detectable in the reporter lines under normal +S growth conditions, GSL levels in the siliques and seeds of *bglu28/30* displayed changes under +S (Figures 2B and 3). The accumulation of all MTX GSL was increased in the siliques of *bglu28/30* (Figure 3). In the seeds of *bglu28/30*, 8-MTO levels were higher than those in the WT (Figure 3).

By contrast, the differences in GSL profile in mature *bglu28/30* plants were more noticeable under −S (Figure 3). In the seeds and siliques of *bglu28/30*, except for the non-detectable 3MSOP levels in the seeds and a slight decrease of I3M level in the siliques, the content of all other GSL species was increased (Figure 3). Similarly, in the flowers of *bglu28/30*, 3MSOP, 4MSOB, 8MSOO, and 4MTB accumulated to a higher level than those in the WT (Figure 3). The increased GSL accumulation in different organs of mature *bglu28/30* suggested that BGLU28 and BGLU30 facilitates GSL catabolism in these organs under S stress.

Surprisingly, the content of 4MSOB, 4MTB, 7MTH, 8MTO, and I3M in the rosette leaves of after-bolting (RLA) *bglu28/30* decreased relative to those in the WT (Figure 3). As plant bolting greatly changes the distribution of GSL in the source and sink organs (Brown et al., 2003; Andersen and Halkier, 2014), we also analyzed the GSL level in the rosette leaves of WT and *bglu28/30* before bolting (RLB). We found that in the 4-week-old before bolting plants, 4MSOB and 4MTB level was still significantly higher in the rosette leaves of *bglu28/30* under −S compared to those in the WT (Figure 3). Notably, 4MTB levels exceeded 4MSOB levels in the mutant under these conditions. These results indicated that the GSL level was more substantially decreased upon bolting in the rosette leaves of *bglu28/30* than in those of WT under −S.

### Source-to-sink transport of GSL upon bolting was induced in *bglu28/30* under −S

Previous studies suggested that GSL is transported from the rosette leaves to reproductive organs upon bolting (Ellerbrock et al., 2007; Andersen et al., 2013; Andersen and Halkier, 2014). Thus, the drastic reduction of GSL level in the rosette leaves of flowering *bglu28/30* could be because of the enhanced GSL transport. To verify this hypothesis, we conducted allyl GSL feeding assay (Figure 4; Andersen et al., 2013). Allyl GSL is a GSL structure that is not produced in *Arabidopsis* Col-0 (Kliebenstein et al., 2001). By feeding allyl GSL to rosette leaves, we can eliminate the interference of endogenous accumulation and biosynthesis.

**Figure 4.**
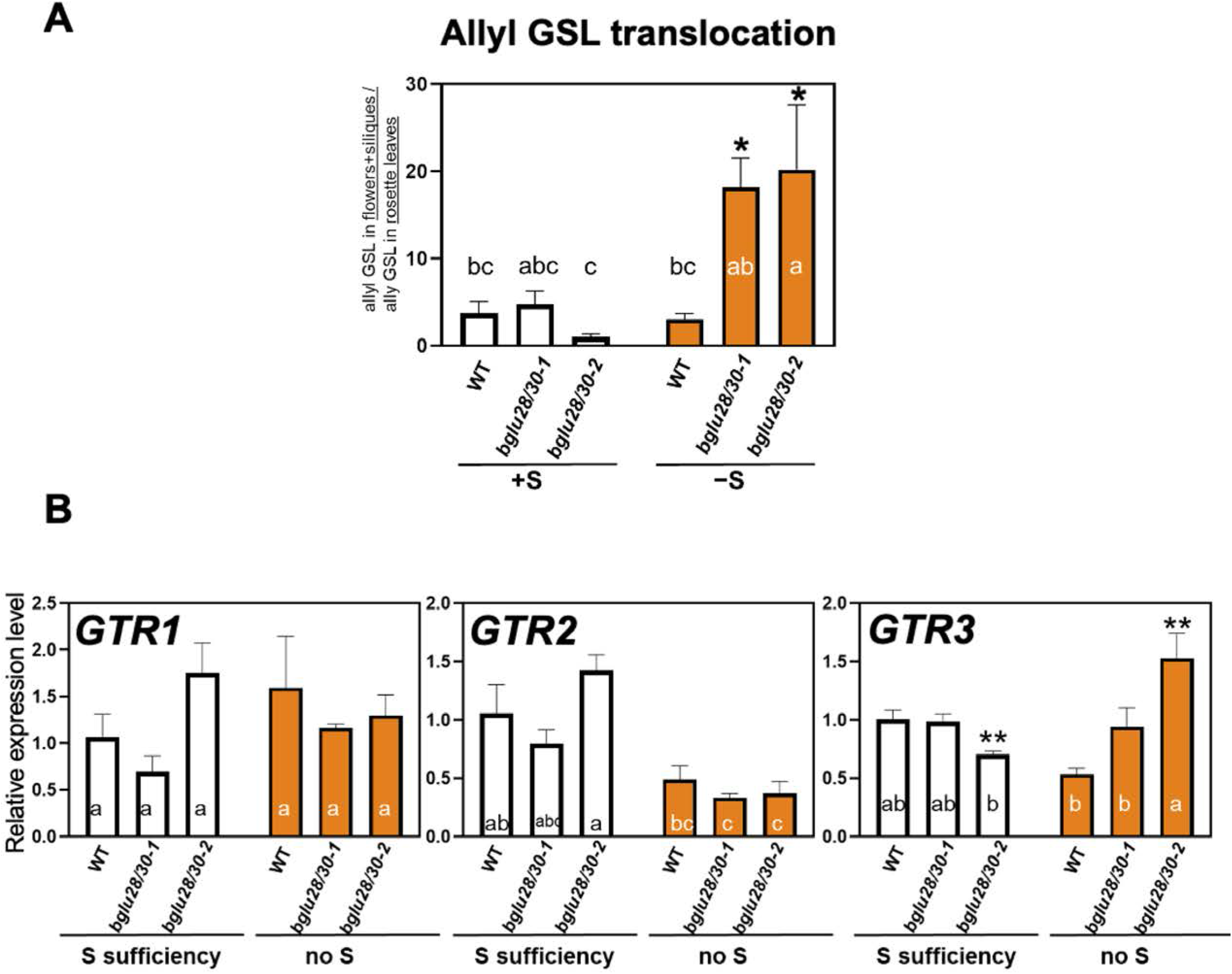
Long-distance transport of rosette leaves-fed allyl GSL in *bglu28/30* and the wild-type (WT) plants grown under different S conditions. (A) Exogenous allyl GSL was fed on the rosette leaves of 5-week-old WT and *bglu28/30* plants grown with sufficient S (+S, white bar) or deficient S (−S, orange bar) for 72 hours. The total content of allyl GSL in the siliques and flowers detected by LC-MS was divided by that in fed-rosette leaves to represent the translocation of allyl GSL between the organs. The values and error bars indicate means ± SEM (n = 4). (B) Relative expression levels of *GTR1, GTR2*, and *GTR3* in the rosette leaves of 5-week-old after bolting *bglu28/30* and WT plants grown under different S conditions (+S, white bar; −S, orange bar) was determined by qRT-PCR. The values and error bars indicate means ± SEM (n = 3). Two-way ANOVA followed by the Tukey–Kramer test was applied to all experimental conditions. Different letters indicate significant differences (*P* < 0.05). Asterisks indicate significant difference detected by one-way ANOVA followed by the Dunnett’s test between *bglu28/30* lines and WT under the same S condition (* 0.05< *P* <0.1, ** 0.01< *P* <0.05).

To rule out potential phenotypes associated with either SALK insertion in our double mutant, we generated an additional *bglu28/30* line (*bglu28/30-2*) and analyzed both *bglu28/30* lines in parallel. The double disruption line used in the former experiments is indicated as *bglu28/30-1* (Figures 1 and 3). In the allyl GSL feeding assay, a total amount of 60 nmol of allyl GSL was fed to rosette leaves of bolted WT and *bglu28/30* plants grown under +S and −S conditions. After 72 hours of treatment, we analyzed the distribution of allyl GSL in the rosette leaves, flowers, and siliques. The distribution ratio of allyl GSL between rosette leaves and reproductive tissues were similar for all genotypes under +S. However, the translocation of allyl GSL to the flowers and siliques was enhanced in *bglu28/30* under S-limited conditions (Figure 4A). These results further supported the idea that GSL transport process in mature plants was stimulated by the malfunction of BGLU28 and BGLU30 under −S.

Translocation of GSL between different organs requires specific transporters. At present, three GSL transporters (GTRs) from the nitrate/peptide transporter (NPF) family have been identified (Nour-Eldin et al., 2012; Andersen et al., 2013; Andersen and Halkier, 2014). Two high-affinity GTRs, GTR1 (NPF2.10) and GTR2 (NPF2.11), are required for the long-distance transport of GSL from sink organs to seeds (Nour-Eldin et al., 2012). GTR3 (NPF2.9) is evolutionarily close to GTR1 and GTR2 and was reported to be an iGSL-specific transporter that probably functions in importing iGSL to S cells in plant roots (Jørgensen et al., 2017). To investigate whether these GSL transporters are responsible for the altered distribution of GSL in the source and sink tissues of *bglu28/30* under −S, we examined their transcript levels in rosette leaves of after-bolting plants (RLA) under different S conditions. The expression level of *GTR1* was neither regulated by S stress nor altered in *bglu28/30*, while that of *GTR2* was downregulated by −S in all genotypes (Figure 4B). Markedly, only the expression level of *GTR3* was increased in the RLA of *bglu28/30* compared to that of WT under −S (Figure 4B). Moreover, *GTR3*’s expression was not affected by the disruption of *BGLU28/30* in plant siliques and before-bolting rosette leaves (RLB) (Supplemental Figure S2). Together, these results implied that GTR3 possibly contributes to the enhanced export of GSL from rosette leaves of after-bolting *bglu28/30* under −S.

### Cysteine and glutathione levels were decreased in the seeds of *bglu28/30* under −S

−S-induced GSL catabolism has been demonstrated to be a growth-sustaining strategy for plant seedlings, by which S released from GSL can be recycled to prioritize the synthesis of primary S-containing metabolites, like cysteine and glutathione (Zhang et al., 2020; Sugiyama et al., 2021). To test whether such optimization of S utilization also happens in mature plants, we then analyzed the contents of cysteine and glutathione in different tissues of WT and *bglu28/30* grown under different S conditions.

Cysteine and glutathione levels in the siliques and rosette leaves before or after bolting of both genotypes were similarly affected by −S; i.e., both cysteine and glutathione in the siliques and RLA were reduced by −S (Figure 5). In RLB, glutathione was decreased, whereas cysteine level was not affected by −S (Figure 5). Noticeably, the accumulation of cysteine and glutathione in the mature seeds of these two genotypes was dissimilar. Even under +S, their levels were either tended or significantly decreased in the *bglu28/30* seeds, and such decrease was more pronounced when S was depleted (Figure 5).

**Figure 5.**
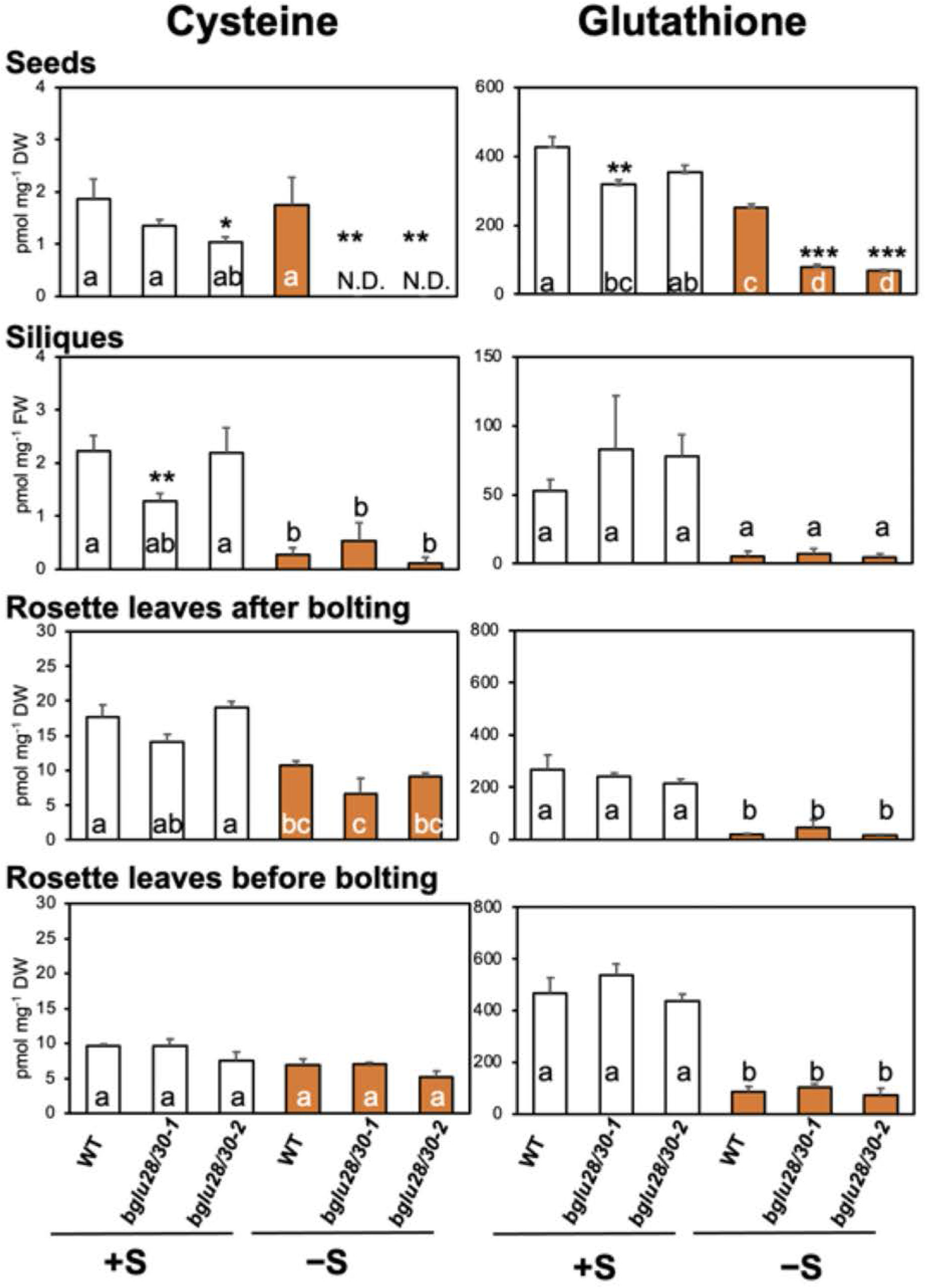
Cysteine and glutathione content in different tissues of *bglu28/30* and the wild-type (WT) plants grown under different S conditions. Plants were grown on vermiculite supplied with sufficient S (+S, white bar) or deficient S (−S, orange bar) for 5 weeks (after bolting) or 4 weeks (before bolting). Cysteine and glutathione content in per mg dry weight (DW) or fresh weight (FW) of each plant tissue was measured using HPLC-fluorescent detection system. The values and error bars indicate means ± SEM (n = 3). Two-way ANOVA followed by the Tukey–Kramer test was applied to all experimental conditions. Different letters indicate significant differences (*P* < 0.05). Asterisks indicate significant difference detected by one-way ANOVA followed by the Dunnett’s test between *bglu28/30* lines and WT under the same S condition (* 0.05< *P* <0.1, ** 0.01< *P* <0.05, *** *P*<0.01).

Because the S-containing amino acid cysteine mainly incorporates into protein, we subsequently examined the S content in the protein fraction of seeds precipitated with 80% ethanol. The total S content in the seeds and both soluble and protein fractions after protein precipitation were decreased by −S in all genotypes (Figure 6). Under −S, the S content in the seeds and the soluble fractions increased in both *bglu28/30* lines (Figure 6). The *bglu28/30* seeds contained ca. 60 nmol S per mg. This difference corresponded well to the higher GSL levels in seeds (Figure 3) taking into account that mGSLs contain three sulfur atoms per molecule. Compared to WT, S content in the seed protein fraction was reduced in the *bglu28/30-2* under +S, but under −S, no change was detected among the three plant lines (Figure 6). Taken together, since BGL*U28/30* disruption reduced accumulation of primary S metabolites and increased levels of GSL in the mature seeds under −S, we concluded that BGLU28/30-dependent GSL catabolism contributes to maintaining the relative levels of primary and specialized S metabolites in seeds.

**Figure 6.**
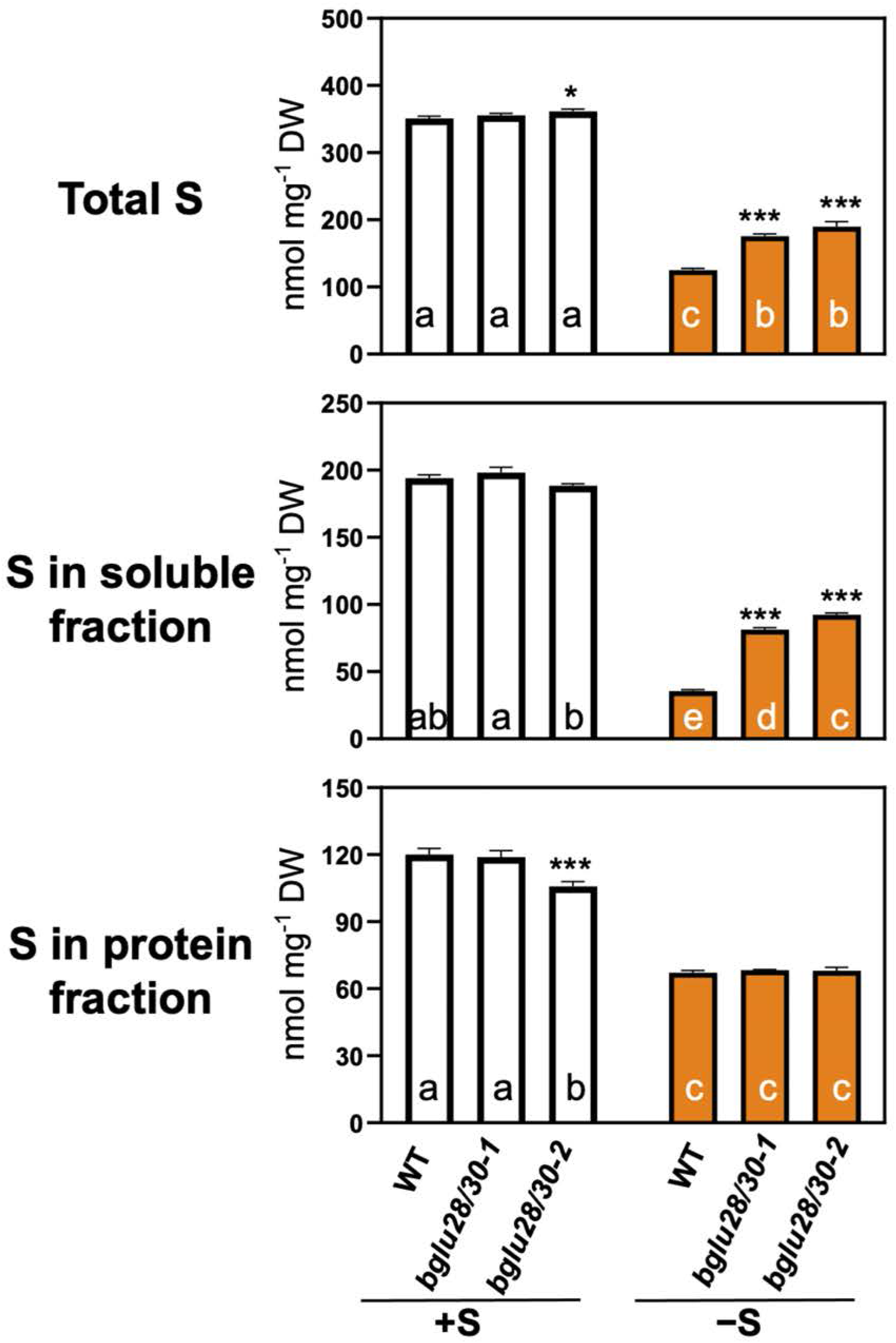
Total S content, S content in soluble and protein fraction of the mature seeds of *bglu28/30* and wild-type (WT) plants grown under different S conditions. Total S content, S content in soluble and protein fraction per mg dry weight (DW) of the mature seeds of *bglu28/30* and the WT plants grown with sufficient S (+S, white bar) or deficient S (−S, orange bar) was determined using ICP-OES. Protein fraction was obtained as the precipitates with 80% ethanol and the supernatants were the soluble fraction. The values and error bars indicate means ± SEM (n = 4). Two-way ANOVA followed by the Tukey– Kramer test was applied to all experimental conditions. Different letters indicate significant differences (*P* < 0.05). Asterisks indicate significant difference detected by one-way ANOVA followed by the Dunnett’s test between *bglu28/30* lines and WT under the same S conditions (* 0.05< *P* <0.1, ** 0.01< *P* <0.05, *** *P*<0.01).

### *bglu28/30* showed an altered transcriptome response under +S and −S

To better understand the effects of *BGLU28/30* disruption on plants, we compared the transcriptome of *bglu28/30* with that of the WT by mRNA-seq using RLA and the developing young siliques grown under different S conditions. The principal component analysis (PCA) was conducted to visualize the difference among the biological groups. For the analysis of silique samples, PC1 (40.5 %), PC2 (19.1 %), and PC3 (13.4 %) contributed to a sum of 73.0 % of the variance. Likewise, 76.6 % of the variance among the RLA samples could be explained by the first three PCs (PC1 48.6 %; PC2 17.4 %; PC3 10.6%) (Supplemental Figure S3). The 3D PCA plot showed that +S and −S samples of both organs were clustered separately, and the samples of different genotypes displayed separation under −S (Supplemental Figure S3). Besides, RLA samples were detached more clearly by different genotypes under both S conditions.

To mine the transcriptional changes brought by *BGLU28/30* disruption, we then examined the differentially expressed genes (DEGs) between WT and *bglu28/30* under both S conditions (Figure 7; Supplemental Table S1). We noticed that, in siliques, there were a larger number of DEGs under −S, while in RLA, the DEGs were more under + S. Furthermore, 5 DEGs in siliques, AT1G75945, *JAL23* (AT2G39330), *AR781* (AT2G26530), *ABCG19* (AT3G55130), *WRKY46* (AT2G46400), and 83 DEGs in RLA, such as *DUF239* (AT2G44240), *CYP81F1* (AT4G37430), *LAX3* (AT1G77690), appeared under both S conditions (Figure 7; Supplemental Table S1).

**Figure 7.**
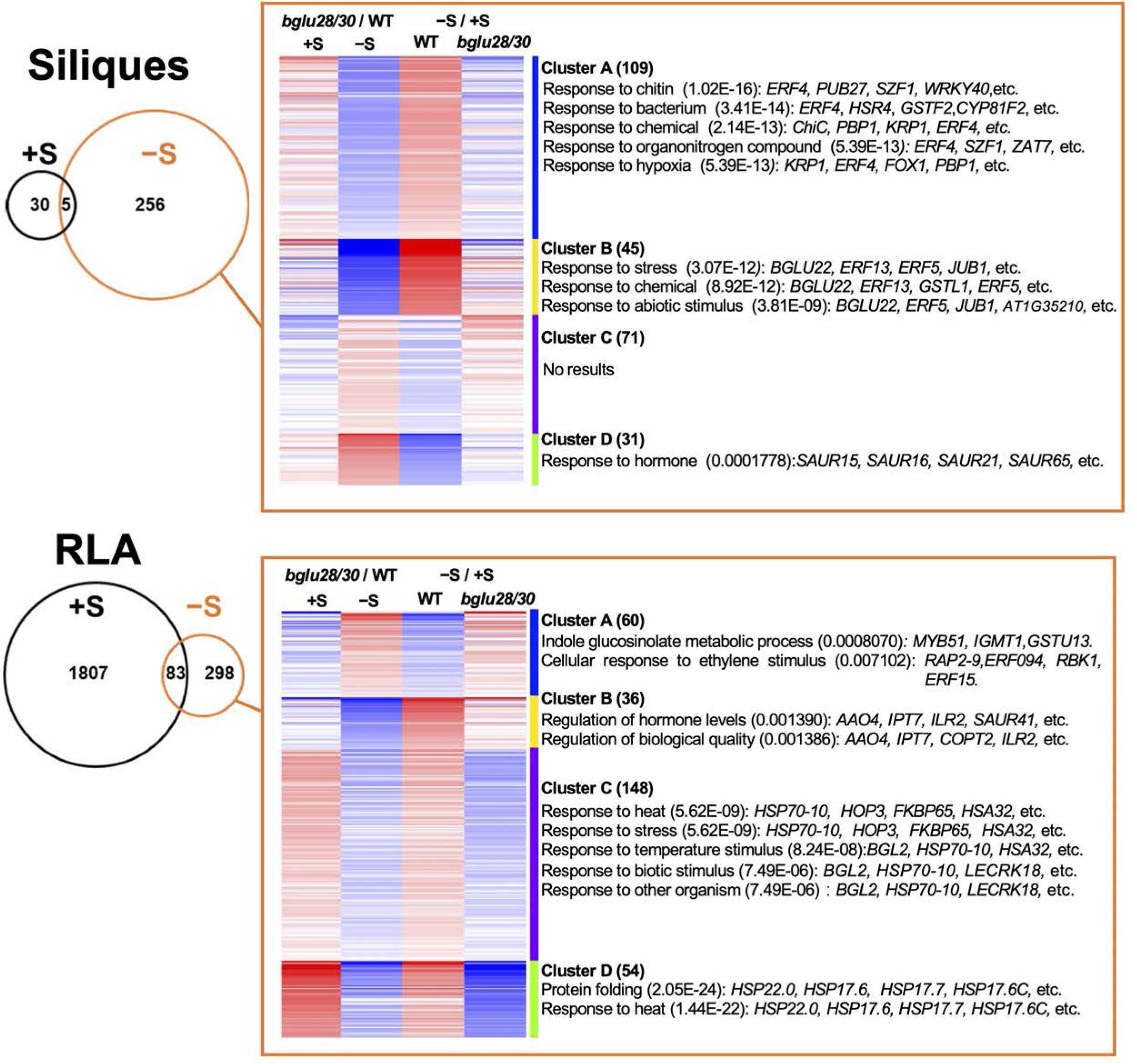
Effects of *BGLU28/30* disruption on the rosette leaf and silique transcriptomes under different S conditions. Venn diagrams of differentially expressed genes (DEGs) in the rosette leaves and siliques of these two genotypes grown under sufficient S (+S, black circle) or deficient S (−S, orange circle) was depicted. Siliques: 7-day after-flowering siliques; RLA: rosette leaves of 5-week-old after bolting plants. Within the orange square: heat map from k-means clustering analysis of DEGs between the genotypes under −S. The fold change value of these DEGs’ expression level between the genotypes under the same S condition, and that between different S conditions in the same genotype were used for the clustering analysis. Red and blue represents a higher and lower expression of the gene in each comparison, respectively. The number of genes in each cluster was shown in the parenthesis. Details about genes included in each cluster is provided in Supplemental Table S3. Enrichment analyses were performed in GO terms of the biological processes category. Representative significantly enriched GO terms (adjusted P values were in the parenthesis) and the genes belonging to the corresponding term are presented.

Because we mainly observed growth and metabolic phenotypes in *bglu28/30* grown under −S, we then focused our investigation on DEGs between the genotypes under −S. To classify these DEGs under −S, we listed the fold change (FC) of their expression level between the genotypes under each S condition and between −S and +S in each genotype, and then conducted K-means clustering on these FC data (Figure 7).

256 DEGs in siliques were divided into four clusters (Figure 7; Supplemental Tables S2, S3). The expression of siliques Cluster A and B genes were less in *bglu28/30* than in the WT under −S. Most Cluster A and B gene expressions were either trending or significantly up-regulated by −S in WT, but their response to −S was compromised in *bglu28/30*. Conversely, the transcript levels of siliques cluster C and D genes showed higher expression in *bglu28/30* than in the WT under −S, and were either trending or significantly down-regulated in WT when S was limited but failed to be repressed by −S in *bglu28/30*. The Gene Ontology (GO) analysis of the genes belonging to each cluster revealed that siliques Cluster A genes were related to plant response to chitin, bacterium, chemical, organonitrogen compound, and hypoxia, represented by the genes such as *ERF4* (AT3G15210), *SZF1* (AT3G55980), *CYP81F2* (AT5G57220), *PBP1* (AT5G54490), *KRP1*(AT4G27280). The

Cluster B genes were enriched to GO terms about plant response to stress, chemicals, and oxygen-containing compounds, represented by *BGLU22* (AT1G66280), *ERF13* (AT2G44840), *ERF5* (AT5G47230), *JUB1* (AT2G43000), *GSTL1* (AT5G02780). As for Cluster C and D genes, Cluster C genes did not show enrichment for GO terms, while Cluster D genes were specifically relevant to plant response to hormone-mediated signaling, especially auxin signaling, as four small auxin-up RNA (SAUR) genes, *SAUR15* (AT4G38850), *SAUR16* (AT4G38860), *SAUR21* (AT5G18030), *SAUR65* (AT1G29460), were sorted to this cluster.

298 DEGs in RLA were also separated into four clusters (Figure 7; Supplemental Tables S2, S3). The expression of RLA Cluster A genes was higher in *bglu28/30* than in the WT under −S, and the suppression of RLA Cluster A gene expression under −S in WT was released or reversed in *bglu28/30*. The expression of RLA Cluster B to D genes was lower in *bglu28/30* than in the WT under −S. Furthermore, these transcripts were principally increased or not changed in WT under −S but could not be identically up-regulated or maintained in *bglu28/30*. The expression of genes belonging to RLA Cluster C and D was even reversely down-regulated by −S in *bglu28/30*. The GO analysis of the four clusters of RLA DEGs exhibited that RLA Cluster A genes, *MYB51* (AT1G18570), *IGMT1* (AT1G21100), *GSTU13* (AT1G27130), were associated with indolic GSL metabolism, and another set of RLA cluster A genes, *ERF15* (AT2G31230), *RBK1* (AT5G10520), *ERF094* (AT1G06160), and *RAP2-9* (AT4G06746), have roles in cellular responses to ethylene. RLA Cluster B genes were involved in regulation of hormone levels, biological qualities, and hormone metabolic process represented by genes like *AAO4* (AT1G04580), *IPT7* (AT3G23630), *ILR2* (AT3G18485), *SAUR41* (AT1G16510), *COPT2* (AT3G46900).

Noticeably, both RLA Cluster C and D genes showed enrichment for plant response to heat due to the intense repression of *heat shock proteins* (*HSPs*) under −S exclusively in *bglu28/30*’s RLA. A list of *HSPs, HSP70-2* (AT5G02490), *HSP70-4* (AT3G12580), *HSP70-5* (AT1G16030), *HSP70-8* (AT2G32120), *HSP70-10* (AT5G09590), *HSP17*.*6A* (AT1G59860), *HSP17*.*7* (AT5G12030), *HSP17*.*8* (AT1G07400), *HSP18*.*1* (AT5G59720), *HSP22*.*0* (AT4G10250), repressed their expression under −S specifically in the *bglu28/30* mutants. Besides, three HSPs, *HSP17*.*6C* (AT1G53540), *HSP17*.*6* (AT5G12020), and *HAS32* (AT4G21320), decreased their expression under −S in RLA of both genotypes, but the reduction was more drastic in the *bglu28/30* mutants.

Overall, the perturbation in GSL catabolism and altered transcriptional responses to S stress in the *bglu28/30* mutants implies crosstalk between GSL metabolism and multiple biological processes under −S.

## Discussion

### BGLU28/30 function in various tissues under −S to catabolize GSL degradation

GSL degradation is stimulated under −S in plant seedlings, and BGLU28/30 were proven to be the main enzymes responsible for this catabolic process (Zhang et al., 2020; Sugiyama et al., 2021). However, whether this is a tissue- and growth stage-specific −S-response has not been answered yet. In this study, we explored this question using mature WT and *bglu28/30* plants grown under different S conditions.

Our results revealed that the disruption of *BGLU28/30* led to increased GSL levels in mature seeds, developing siliques, flowers, and RLB under −S (Figure 3). Since the transcript level of most GSL biosynthetic genes was not increased in the tissues of *bglu28/30* compared to those in the WT under −S, the accumulation of GSL in these organs of *bglu28/30* was likely not caused by enhanced GSL biosynthesis (Supplemental Table S4). Together with the observation that −S strongly induced the expression of *BGLU28/30* in these organs, we concluded that BGLU28/30 might catabolize GSL in specific organs of mature plants in response to −S (Figure 2). Considering the presence of S cells which store a GSL at high concentrations in these reproductive organs and leaves, we propose that expression of *BGLU28/30* in these organs under −S contributes to the efficient recycling of S released from GSL (Koroleva et al., 2000, 2010; Sarsby et al., 2012). Moreover, as the mature *bglu28/30* displayed reduced biomass and seed yield under −S, S recycling from GSL is required for plant reproductive growth when S is limited (Figure 1).

### Disruption of *BGLU28/30* stimulates GSL translocation under −S

Apart from GSL catabolism, we further found that when *BGLU28/30* were disrupted, GSL distribution towards the siliques and flowers of plants grown under −S was stimulated (Figure 4A). There could be several possible explanations for this phenotype. For instance, plants may be able to sense S status in developing seeds and transduce an unknown signal to stimulate the GSL translocation. Alternatively, the reduced levels of GSL catabolites in *bglu28/30* RLA tissue might be an indirect cause that affected GSL translocation. This possibility was inspired by our transcriptome analysis which is discussed later.

Moreover, the increased expression of *GTR3* suggests that altered GTR activity may result in enhanced GSL translocation in *bglu28/30* under −S (Figure 4B). Furthermore, expression of *GTR3* in phloem companion cells supports it role in counteracting long-distance phloem transport (Wang and Tsaya, 2011). Nevertheless, GTR3 is reported to prefer iGSL, but we did not observe an apparent increase of iGSL in the flowers and siliques of *bglu28/30* grown under −S (Figure 3). In *Arabidopsis* aerial parts, mGSL are generally more abundant than iGSL and GTR3 was still able to uptake mGSL when iGSL was absent or at a low concentration (Jørgensen et al., 2017). To test the potential contribution of GTR3 in this phenotype, introducing the loss-of-function *GTR3* mutation into the *bglu28/30* mutants would be helpful. Additionally, although transcript levels of *GTR1/2* in *bglu28/30* were similar to that in WT, cell-specific differential expression might have escaped the analysis and/or posttranslational regulation of GTR1/2 might be altered in *bglu28/30*, leading to increase *in planta* transport activity in the mutant.

### Primary metabolites levels decreased in the mature seeds of *bglu28/30* mutants under −S

Under −S, the accumulation of primary S metabolites, cysteine and glutathione, were decreased in the mature plant organs (Figure 5; Morikawa-Ichinose et al., 2019). Although GSL catabolism are thought to support primary S metabolism when S is insufficient (Zhang et al.,2020; Sugiyama et al.,2021), in comparison between WT and *bglu28/30*, levels of these primary S metabolites were indistinguishable between the genotypes in their siliques, RLA and RLB under −S (Figure 5). Our mRNA-seq data also revealed that the transcript level of S assimilatory genes in the silique and RLA of these two genotypes were similar, suggesting that the primary S metabolism was not differentially regulated between the genotypes in these organs (Supplemental Table S4). The effect of *BGLU28/30* disruption on the accumulation of cysteine and glutathione under −S was specific to the mature seeds (Figure 5), indicating that the levels of these primary S metabolites cannot be maintained in seeds in the absence of functional BGLU28/30.

We noticed that seeds distinctively accumulated a high level of 4MTB, while in the siliques and other tissues, 4MSOB is the dominant one (Brown et al., 2003; Meier et al., 2019; Figure 3). Our previous report revealed that the accumulation of 4MTB was most drastically affected by *BGLU28/30* disruption in plant seedlings grown under −S (Zhang et al., 2020). Indeed, BGLU28 exhibited a higher hydrolysis rate against 4MTB than other GSL species (Sugiyama et al., 2021). Since mature seeds accumulate a substantial amount of 4MTB and BGLU28/30 prefer to hydrolyze this GSL species, more S can be released from GSL by the activity of these two BGLUs during seed development and eventually incorporated into the primary metabolite under −S. Therefore, loss-of-function BGLU28/30 have more profound impact on the primary S metabolite accumulation in seeds. Based on these findings, we propose that BGLU28/30 catalyze a key step in the allocation of primary and specialized S metabolites in the seeds responding to the S status in the plant.

Besides the above discussion, two more questions deserve attention. One is why *Arabidopsis* (Col-0) seeds mainly accumulate 4MTB. The conversion of 4MTB to 4MSOB is catalyzed by the five flavin-monooxygenases, FMO GS-OX1 to 5 in *Arabidopsi*s (Li et al., 2008, 2011). Expression of *FMO GS-OX1-5* was not found in seeds, but most 4MTB accumulated in seeds was converted to 4MSOB by the artificial overexpression of *FMO GS-OX1* (Hansen et al., 2007; Li et al., 2011). Thus, lacking an active FMO GS-OX in seeds may cause this special seed GSL profiles. Besides, as discussed in a previous review (Nour-Eldin and Halkier, 2008), unidentified enzymes which catalyze the conversion of 4MSOB to 4MTB could exist in seeds or plants may selectively transport 4MTB from siliques to seeds. However, the physiological relevance of such seed-specific GSL profile is still unclear. Also, why 4MTB is a favored substrate for BGLU28/30 under S stress is a question that needs an answer.

### Transcriptional responses to S stress were changed in the *bglu28/30* mutants

Our transcriptome analysis disclosed that *BGLU28/30* disruption unexpectedly altered plant responses to −S. In the RLA of WT, −S repressed the expression of ethylene response-related genes, *ERF15, RBK1, ERF094*, and *RAP2-9*, but not in *bglu28/30* (Figure 7; Supplemental Tables S2 and S3). In the siliques, the expression of ethylene response factors, *ERF13, ERF5*, and *ERF4*, also failed to be activated by −S in *bglu28/30* as it was observed in the WT (Figure 7; Supplemental Table S2 and S3). In the RLA of WT, −S repressed the expression of ethylene response-related genes, *ERF15, RBK1, ERF094*, and *RAP2-9*, but not in *bglu28/30* (Figure 7; Supplemental Tables S2 and S3). In the silique, the expression of ethylene response factors, *ERF13, ERF5*, and *ERF4*, also failed to be activated by −S in *bglu28/30* as in the WT (Figure 7; Supplemental Table S2 and S3).

Ethylene production was shown to increase in response to −S and this phytohormone is proposed to be a signaling molecule that adjusts S metabolism under S limited conditions (Moniuszko et al., 2013; Wawrzynska et al., 2015). A recent paper reported that exogenous ethylene treatment led to a rapid reduction of GSL levels, further unveiled the role of ethylene in regulating accumulation of these specialized S-containing metabolites (Hildreth et al., 2020). In agreement with the earlier findings and hypotheses, our observations indicate that ethylene is involved in plant responses to S limitation in above ground tissues. Additionally, GSL and/or derived metabolites may reversely affect ethylene production/signaling under this nutrient stress.

Besides ethylene-related genes, the auxin signaling-related genes, *SAUR15, SAUR16, SAUR21*, and *SAUR65*, were downregulated under –S in the siliques of the WT, but not in *bglu28/30* (Figure 7; Supplemental Tables S2 and S3). Similarly reduced *SAUR* gene expression was previously observed under other abiotic stresses, for example osmotic stresses by the transcriptional repressors, *Arabidopsis* zinc-finger protein 1 (AZF1) and AZF2 (Kodaira et al., 2011). It is unclear how BGLU28/30-mediated GSL catabolism may contribute to the repression of *SAURs* under –S. Although GSL catabolism could be associated with auxin signaling (Katz et al., 2020), since other hormones also regulate *SAUR* expression, we cannot affirm that their altered expression under −S and in *bglu28/30* is linked to the perturbation of auxin level (Ren and Gray, 2015; Stortenbeker and Bemer, 2019). The mechanism responsible for the downregulation of *SAURs* under −S requires further studies.

In addition to the phytohormone-related genes that showed altered responsiveness to −S in *bglu28/30*, another noteworthy transcriptional pattern in these mutants is the pronounced repression of *heat shock proteins* (*HSPs*) under −S in RLA (Figure 7; Supplemental Tables S2 and S3). This observation indicated that GSL or their breakdown products may be involved in modulating the expression of *HSPs*. In line with our speculation, previous reports found that one class of GSL breakdown products, isothiocyanate, could induce the expression of *HSPs* and a GSL-deficient line was unable to accumulate *HSPs* upon heat stress (Ludwig-Müller et al., 2000; Hara et al., 2013). The reduced expression of *HSPs* might explain the growth and metabolic changes in *bglu28/30* under −S. HSPs are classified into different groups based on their approximate molecular weight (Al-Whaibi, 2011). The *HSPs* downregulated under –S in the *bglu28/30* are mainly from the HSP70 family, and the small HSP family (Supplemental Tables S2 and S3). Both HSP70 and small HSPs are proposed to act as chaperones to sustain cell hemostasis under stressed or non-stressed conditions (Sung et al., 2001; Waters and Vierling, 2020). Dysfunction of certain HSP70 and small HSPs members resulted to retarded plant growth (Su and Li, 2008; Aghaie and Tafreshi, 2020; Escobar et al., 2021). Thus, the reduced expression of these HSPs could result in the reduced growth observed for *bglu28/30* under −S. Furthermore, the loss of HSP70 and small HSPs members can damage cell membrane integrity (Aghaie and Tafreshi, 2020; Escobar et al., 2021). So, if the low expression of these *HSPs* in *bglu28/30* under −S led to negatively affected cell membrane integrity, GSL stored in the RLA cells of *bglu28/30* could be leaked and finally loaded to phloem, which caused the observation that GSL transport was enhanced in these mutants. Besides, the reduced expression of *HPSs* is likely to indicate the change of intracellular pH, which may increase the proton gradient-driven activity of the GTRs (Triandafillou et al., 2020; Chung et al., 2022). To clarify these possibilities, the contribution of HSPs to the phenotypes of *bglu28/30* under −S needs to be investigated.

Lastly, disruption of *BGLU28/30* caused transcriptional changes in many genes also when S was sufficient (Figure 7; Supplemental Table S1). Coupled with the result that the siliques of *bglu28/30* accumulate more GSL than that of the WT plants under +S, we propose that these two BGLUs also function during plant development in the absence nutritional stress (Figure 3).

## Conclusion

In summary, our study confirmed the role of BGLU28/30 in catalyzing GSL catabolism in rosette leaves and reproductive organs to sustain plant growth under −S. The induced GSL transport from rosette leaves to the reproductive organs in *bglu28/30* under −S implies an interconnection between GSL catabolism and distribution at the whole plant level. The differential expression of genes related to phytohormone signaling and heat responses in *bglu28/30* under −S further supports the need for crosstalk between GSL catabolism and these biological processes. Overall, these findings broadened our knowledge about the function of these two atypical BGLUs and the physiological significance of GSL catabolism under S stress.

## Materials and Methods

### Plant Materials and Growth Conditions

The *Arabidopsis thaliana* accession Columbia (Col-0) was used as the wild-type (WT). The *bglu28/30-1* mutants carry T-DNA insertion in the fourth exon of *BGLU28* (At2g44460, SALK_043339) and the eighth exon of *BGLU30* (At3g60140, SAIL_694_G08) (Zhang et al. 2020). *bglu28/30-2* carry T-DNA insertion in the third intron of *BGLU28* (SALK_011332) and the tenth intron of *BGLU30* (SAIL_861_F02). Both double insertion lines were generated by the cross-fertilization of the single insertion lines obtained from the Arabidopsis Biological Resource Center (ABRC). Homozygosity of all insertions was confirmed at the gene and the transcript levels by genotyping PCR and RT-PCR, respectively, as described previously (Zhang et al., 2020).

For the seedling growth, plants were grown on MGRL agar medium containing 1% sucrose and 0.5% agarose, at 22 °C under constant illumination (40 µmol m^−2^ s^−1^) (Fujiwara et al., 1992; Hirai et al., 1995). Agar medium was prepared as described previously (Kimura et al., 2019). Sufficient S (+S) MGRL agar medium was supplemented with 1500 µM MgSO_4_. Deficient S (–S) MGRL agar medium did not contain MgSO_4_ and 1500 µM MgCl_2_ was supplied to adjust Mg concentration.

For mature plant growth, plants were cultured in a growth chamber controlled at 23 ± 2 °C under constant illumination (40 µmol m^−2^ s^−1^). The seeds of WT, *bglu28/30-1*, and *bglu28/30-2* were sown in 5 × 5 × 5 cm plastic pots filled with vermiculite. MGRL solution containing 750 µM MgSO_4_ (+S, sufficient S) or 0 µM MgSO_4_ (–S, deficient S) was supplied to the plants twice a week. For –S conditions, Mg^2+^ concentration in the solution was adjusted to 750 µM with MgCl_2_. Five weeks after sowing the seeds, different plant tissues, rosette leaves, stems, flowers, and siliques were harvested separately from each pot. Rosette leaves were weighed for the fresh weights. Primary stem height was measured six weeks after seed sowing. Seeds were collected and weighed when plants were utterly mature, and the remaining parts after seed harvesting were dried and measured for the dry weight of the aerial part. The harvested plant tissues and seeds were then frozen in liquid nitrogen, freeze-dried, ground into a fine powder using a Tissue Lyser (Retsch, Haan, Germany), and used for the subsequent metabolites analysis.

### Observation of *BGLU28* and *BGLU30* expression using promoter-YFP lines

5’upstream regions of *BGLU28* (*P*_*BGLU28*_, −2928 bp from translational start codon) and that of *BGLU30* (*P*_*BGLU30*_, −2045 bp from translational start codon) were amplified with the primer pairs listed in Supplemental Table S5. The PCR-amplified *P*_*BGLU28*_ was directly cloned into pENTR/D-TOPO (Invitrogen, USA) and *P*_*BGLU30*_ was cloned into pENTR/D-TOPO digested with the restriction enzymes, AscI and NotI (Takara Bio, Japan). After confirming their sequences, *P*_*BGLU28*_ and *P*_*BGLU30*_ were then subcloned into pMpGWB107 (Ishizaki et al., 2015) using LR Clonase II (Life Technologies, USA). The recombinant plasmids were transformed to *Agrobacterium* and then integrated into the WT plants by floral dip transformation (Clough and Bent, 1998). In the *T*_*1*_ generation, transgenic plants were selected on the GM media containing 50 µg mL^−1^ of carbenicillin and hygromycin. Selected plants were grown on the soil to obtain the *T*_*2*_ generation (*P*_*BGLU28*_-YFP and *P*_*BGLU30*_-YFP).

To observe the yellow fluorescence derived from YFP in seedlings, both transgenic and the WT plants were grown on MGRL medium containing +S or −S for 7 days and yellow fluorescence was observed with a fluorescent microscope system (EVOS FL Auto 2 Imaging System) equipped with the EVOS Light Cube, YFP (Excitation: 500/24, Emission: 542/27) (Thermo Fisher Scientific, USA).

To observe the YFP fluorescence in mature plants, both transgenic and the WT plants were grown on the GM media for 2 weeks and grown on the vermiculite supplied with +S or −S for another 3 weeks. The yellow fluorescence and the autofluorescence derived from chlorophyll were visualized by an image analysis scanner (Amersham Typhoon, RGB, Cytiva, USA) with a 488 nm excitation laser and Cy2 525BP20 and Cy5 670BP30 emission filters, respectively.

### Glucosinolate analysis

Three mg of plant tissues powder was extracted with 300 µL of ice-cold 80% methanol containing 2 µM L (+)-10-camphor sulfonic acid (10CS, internal standard for negative ionization mode, Tokyo Kasei, Japan). After homogenization, cell debris was removed by centrifugation (13,000 rpm, 15 min, 4 °C), and the supernatants were evaporated with vacuum concentrator (SpeedVac SPD1030, Thermo Fisher Scientific, USA). Dried supernatants were dissolved into 100 µL ultra-pure water and filtrated with 0.2 µM SY4VF (PVDF) syringe filter (Advanced Microdevices Pvt. Ltd., India) prior to the analysis.

GSL levels were analyzed using a high-performance liquid chromatography connected to a triple quadrupole-MS (LCMS8050, Shimadzu, Kyoto, Japan) using L-column 2 ODS (pore size 3 µm, 2.1 × 150 mm, CERI, Japan) as described previously (Morikawa-Ichinose et al., 2019). The mobile phases were as follows: A, ultra-pure water with 0.1% formic acid; B, acetonitrile with 0.1% formic acid. The gradient program was as follows with a flow rate of 0.3 mL/min, 0–0.1 min, 1% B; 0.1–15.5 min, 1-99.5% B; 15.5–17 min, 99.5% B; and 17.1– 20 min, 1% B. The mass spectrometer was run in negative electrospray mode. The precursor ion [m/z]/product ion [m/z] scores of each GSL are shown as follows: 3MSOP: 422.02/358.02, 422.02/96.9, 422.02/95.9; 4MSOB: 436.05/96.9, 436.05/96.0, 436.05/177.9; 8MSOO: 492.1/428.1, 492.1/96.9, 492.1/95.9; 4MTB: 420.04/96.9, 420.04/95.9, 420.04/74.9; 7MTH: 462.09/96.9, 462.09/95.9, 462.09/74.9; 8MTO: 476.11/96.9, 476.11/95.9, 476.11/74.9; I3M: 447.05/96.9, 447.05/95.9, 447.05/74.9; Allyl GSL: 358.05/75, 358.05/96.95, 358.05/95.9. The common product ion 96.9 m/z was used for quantification in all cases. GSLs levels were quantified based on calibration curves using standard GSL compounds, 4MTB, 3MSOP, 4MSOB, and I3M (Cfm Oskar Tropitzsch GmbH, Marktredwitz, Germany) and Allyl GSL (Tokyo Chemical Industry Co., Ltd, Japan).

### Allyl glucosinolate feeding assay

GSL transport from rosette leaves to the reproductive tissues was analyzed as described previously with minor modification (Andersen et al., 2013; Andersen and Halkier, 2014). In brief, allyl GSL was dissolved in –S MGRL solution to a final concentration of 10 mM. 6 µl of it was fed on the rosette leaves of 5-week-old WT and *bglu28/30* plants grown under +S and – S. After 72 hours of feeding, the rosette leaf blades were rinsed with deionized water and then harvested separately from flowers and siliques. The harvested plant tissues were frozen in liquid nitrogen and grind to a fine powder using a Tissue Lyser (Retsch, Germany). Allyl GSL levels in these plant tissues were quantified using LCMS as described in the “Glucosinolate analysis” section.

### Cysteine and glutathione analysis

The plant extracts used for GSL analysis were used for the analysis of cysteine and glutathione. Cysteine and glutathione levels were determined by HPLC-fluorescent detection system (JASCO, Japan) after labeling of thiol bases with monobromobimane (Invitrogen, USA) as described previously (Kimura et al., 2019). The labeled products were separated by HPLC using the TSKgel ODS-120T column (150 × 4.6 mm, TOSOH, Japan) and detected with a fluorescence detector FP-920 (JASCO, Japan) by monitoring the fluorescence at 478 nm under the excitation at 390 nm. The elution was performed with solvent A (12 % methanol, 0.25 % acetic acid) and solvent B (90 % methanol, 0.25 % acetic acid) under the gradient program from 0 % to 25 % of solvent B. Cysteine and glutathione (Nacalai Tesque, Japan) were used as standards.

### Total S analysis

Five mg of mature seed powder was digested in 200 µL of conc. HNO_3_ (KANTO CHEMICAL, Japan) at 95 °C for 30 min followed by an incubation at 115 °C for 90 min. After confirming complete digestion, the samples were cooled to room temperature, diluted to 1 mL with ultra-pure water, and filtrated with 0.45 µm filters (DISMIC-03CP, ADVANTEC, Japan). The processed samples were diluted 5-fold in 0.1 M HNO_3_ containing gallium (KANTO CHEMICAL, Japan) as an internal standard immediately before the analysis. We analyzed S levels using inductively coupled plasma optical emission spectrometer (ICP-OES, Agilent 5800, Agilent Technologies, CA, USA). S levels were quantified based on the standard curve derived from the sequential dilution of S standard solution (KANTO CHEMICAL, Japan). The emission spectrum of S was collected with 180.669 nm.

### S in protein and soluble fraction analysis

Five mg of mature seed powder was homogenized with 300 µL of 80 % ethanol and placed under 4 °C to precipitate protein as described previously (Zhang et al., 2020). The precipitates (protein fraction) and the collected supernatants (soluble fraction) were dried by vacuum evaporation using a vacuum concentrator (SpeedVac SPD1030, Thermo Fisher Scientific, USA). Dried supernatants and precipitates were digested with conc. HNO_3_ (KANTO CHEMICAL, Japan), analyzed using ICP-OES, and S content was quantified as described in the former section, “Total S analysis”.

### mRNA-seq analysis

Two RNA samples from rosette leaves of after-bolting WT plants and *bglu28/30-2* mutant plant grown under different S conditions and their 7-day-post-anthesis siliques were extracted with the RNeasy Plant Mini Kit (QIAGEN, Germany). The residual DNA was removed with DNase treatment (DNaseI, NIPPON GENE, Japan) and total RNA was purified with the SPRI magnetic beads (RNAClean XP, Beckman Coulter Life Sciences, USA). The quantity and quality of RNA samples was determined using MultiNA (Shimazu, Japan).

The isolation of poly-A mRNA and construction of cDNA libraries for strand-specific RNA-Seq were performed using the NEB Next Poly(A) mRNA Magnetic Isolation Module and NEB Next Ultra II Directional RNA Library Prep Kit for Illumina with NEB Next Multiplex Oligos for Illumina (New England Biolabs Inc., USA), respectively. Libraries were subjected to 150-bp paired end sequencing on the HiSeq X platform (Illumina Inc., USA). Raw sequence data were cleaned using Trimmomatic (version 0.39) with the following default parameter settings: ILLUMINACLIP: adapters/TruSeq3-PE.fa: 2:30:10, LEADING: 3, TRAILING: 3, SLIDINGWINDOW: 4:15, and MINLEN: 36. The RNA-Seq reads were mapped to the reference genome of *A. thaliana* (TAIR10) using HISAT2 (version 2.2.1) with the option “--rna-strandness RF” for mapping reverse complement paired-end reads. The quantification of gene expression level was performed using HTSeq (version 0.13.5). Normalization of gene expression data across the libraries and statistical analysis of differential gene expression were performed using edgeR (version 3.38.4). PCA was performed using the “precomp” function in R (version 3.4.1), and the PCA plot was created in Microsoft Excel with the XLSTAT 2022.1 add-in. RNA-seq data has been deposited in National Center for Biotechnology Information (NCBI) Sequence Read Archive (SRA) with accession numbers xxxxx.

### Selection and analysis of the genes differently expressed between S conditions and genotypes

The genes differentially expressed between the genotypes or S conditions were selected with the following parameters: –S versus +S, *bglu28/30* versus WT; log2|Fold Change| ≥1, and q___edgeR value ≤ 0.05. DeepVenn (https://www.deepvenn.com/; Hulsen, 2022.) was used to draw the venn diagrams and g: Profiler (https://biit.cs.ut.ee/gprofiler/gost; Raudvere et al., 2019)was used to perform the Gene onthology (GO) analysis. iDep.96 (http://bioinformatics.sdstate.edu/idep/; Ge et al., 2018) was used for K-means clustering analysis and the following GO enrichment analysis for each cluster.

### qRT-PCR analysis

Reverse transcription was conducted using PrimeScript RT Reagent Kit with gDNA Eraser (Takara, Japan). Real-time PCR was carried out using a KAPA SYBR FAST qPCR Master Mix 2X (Kapa Biosystems, USA), a qTOWER3 G touch (Analytik Jena AG, Germany), and gene-specific primers listed in Supplemental Table S5. Primer efficiency of each primer set was assessed before the experiment. The relative mRNA abundances were calculated with ΔΔCt methods using *UBQ2* as an internal control.

### Statistical analysis

Two-way ANOVA followed by the Tukey–Kramer test and One-way ANOVA followed by the Dunnett’s test were performed using GraphPad Prism9.4.1 (GraphPad Software, USA, https://www.graphpad.com/). Significant differences detected by the Tukey–Kramer test were shown with different letters (*P* < 0.05). Significant differences detected by the Dunnett’s test were shown with asterisks (* 0.05 < *P* <0.1, ** 0.01 < *P* < 0.05, *** *P*<0.01). Student’s *t*-test were carried out in Excel and significant differences are shown with asterisks (* 0.05< *P* <0.1, ** 0.01< *P* < 0.05, *** *P*<0.01).

## Supplemental Data

**Supplemental Figure S1**. No YFP fluorescence was observed in WT plants grown under different S conditions.

**Supplemental Figure S2**. Relative expression level of *GTR3* in RLB and siliques of WT and *bglu28/30* under different S conditions.

**Supplemental Figure S3**. Principal component analysis (PCA) of transcriptome of WT and *bglu28/30* under different S conditions.

**Supplemental Table S1**. DEGs between WT and *bglu28/30* under different S conditions.

**Supplemental Table S2**. −S-responsive genes in WT and *bglu28/30*.

**Supplemental Table S3**. K-means clustering and GO enrichment analysis of DEGs between WT and *bglu28/30* under −S.

**Supplemental Table S4**. mRNA_seq results about GSL metabolism- and S assimilation-related genes’ expression in WT and *bglu28/30* under different S conditions.

**Supplemental Table S5**. Primers used in this study.

## Acknowledgements

We gratefully acknowledge the Arabidopsis Biological Resource Center (ABRC) for providing the T-DNA insertion lines of *BGLU28* and *BGLU30*. The ICP-OES (Agilent 5800) and LCMS-8050 analysis were performed at the Center of Advanced Instrumental and Educational Supports, Faculty of Agriculture, Kyushu University, with the kind instruction by Emiko Matsunaga and Taiki Akasaka, respectively. Plant growth and seed harvesting were done at Biotron Application Center, Kyushu University.

