## Supplemental Figures for "Glucosinolate catabolism maintains glucosinolate profiles and transport in sulfur-starved *Arabidopsis*"

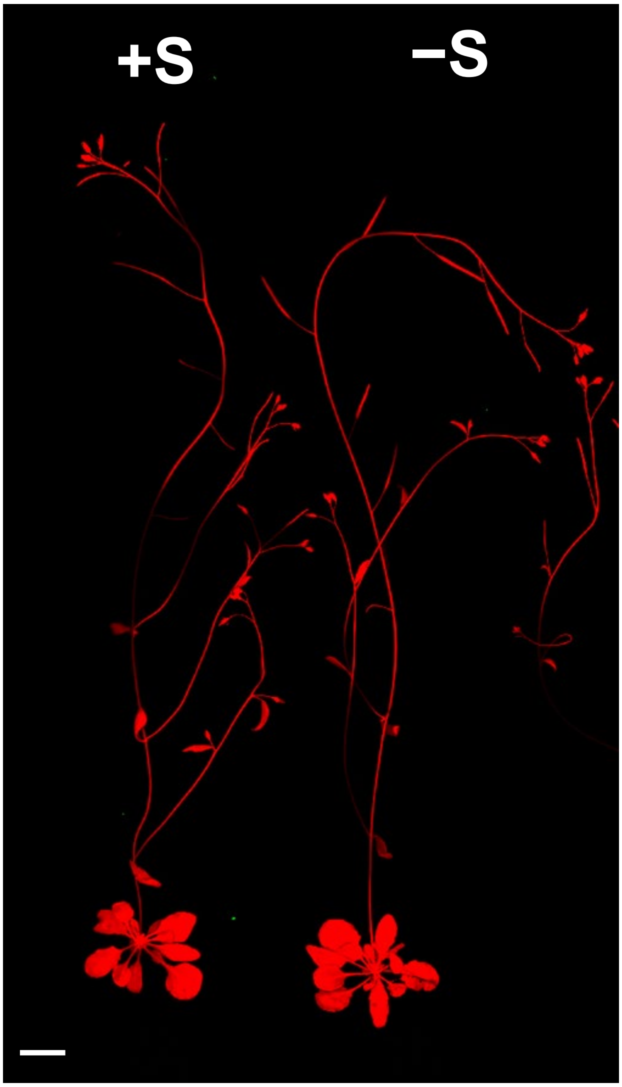


**Supplemental Figure S1. No YFP fluorescence was observed in WT plants grown under different S conditions**

The representative image of 5-week-old WT plants obtained by an image analysis scanner. WT plants were grown together with $P_{BGLU28}$-*YFP* and $P_{BGLU30}$-*YFP* (Figure 2) lines under both +S and −S conditions. The yellow fluorescence and the autofluorescence derived from chlorophyll of WT plants and transgenic lines were monitored simultaneously.

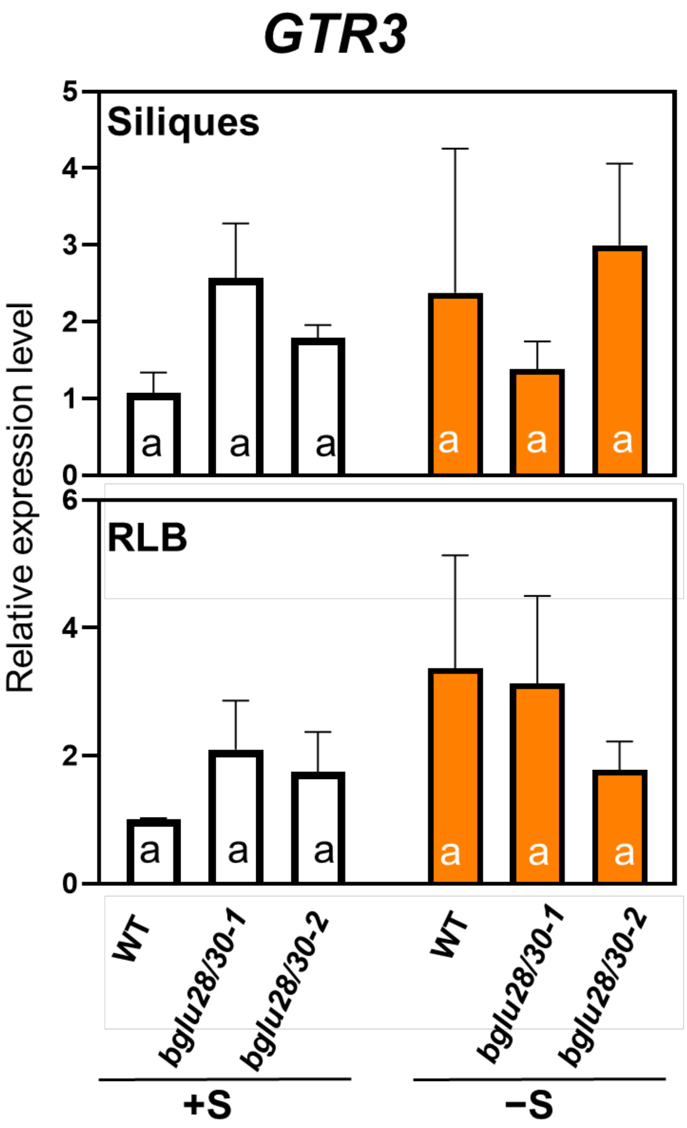


**Supplemental Figure S2. Relative expression level of *GTR3* in RLB and siliques of WT and *bglu28/30* under different S conditions.** Relative expression level of *GTR3* in the rosette leaves of 4-week-old before-bolting (RLB) WT and *bglu28/30*, and their 7-day after-flowering siliques under different S conditions (+S, white bar; −S, orange bar) was determined by qRT-PCR. The values and error bars indicate means ± SEM (n = 3). Two-way ANOVA followed by the Tukey–Kramer test was applied to all experimental conditions. Different letters indicate significant differences ($P < 0.05$). One-way ANOVA followed by the Dunnett's test between *bglu28/30* mutant lines and WT under the same S condition was conducted but no significant differences were detected.

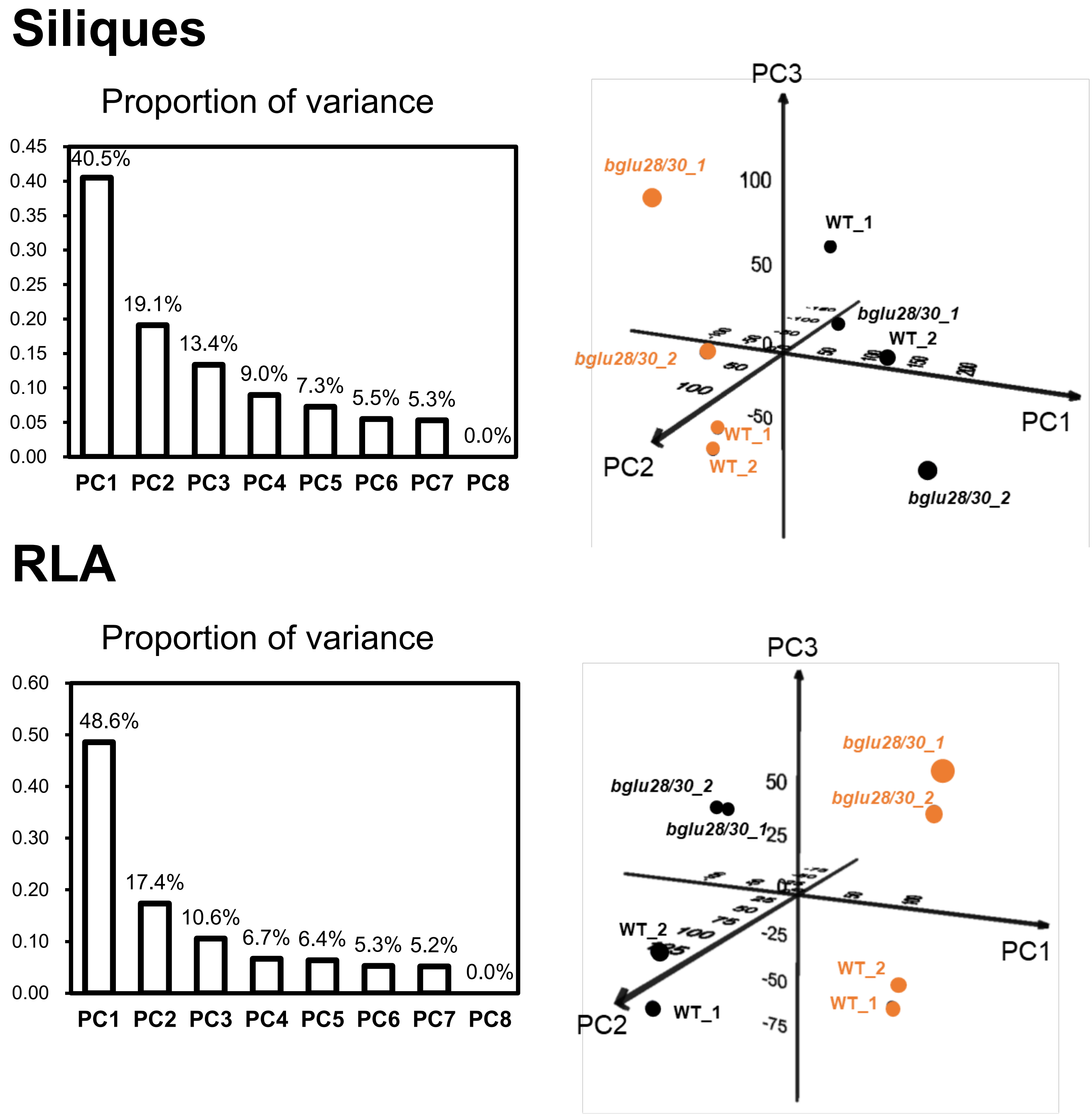


**Supplemental Figure S3. Principal component analysis (PCA) of transcriptome of WT and *bglu28/30* under different S conditions.** The transcriptome data of duplicate samples of 7-day after-flowering siliques, and rosette leaves of 5-week-old after-bolting (RLA) WT and *bglu28/30* (*bglu28/30-2* lines) grown under different S conditions was used for PCA analyses. The proportion of variance explained by each PC is marked on the bar graphs. In PCA plots, black and orange dots indicate samples from plants grown under +S and −S, respectively.
