## Supplemental Table S5 for "Glucosinolate catabolism maintains glucosinolate profiles and transport in sulfur-starved *Arabidopsis*"

**Table S5. Primers used in this study**

| | |
|---|---|
| **Amplification of promoter regions upstream of *BGLU28* and *BGLU30* (5' to 3')** | |
| BGLU28_ups_F | caCCAATGTGAAGGGATTTTTCGCATGGTCGTTGCTA |
| BGLU28_ups_R | ATTCCTTTTTTGTGTTTTGGAATTTAAAAGTGTTATGTATTTTCTTCAAACTGT |
| BGLU30_ups_F | caaaaaagcaggctccgcTATTAGTTTTGTCCCAAACGTTAAACC |
| BGLU30_ups_R | caagaaagctgggtcggTTTTCTCTCTCTCTCTCTCTCTTTTTTTTTT |
| **qRT-PCR (5' to 3')** | |
| GTR1_qF | GCAGCCTTGGGTCAATCTTTACAAC |
| GTR1_qR | GGGGGTCATAATTGCTGCTTTGTC |
| GTR2_qF | ACAAGCAGTTTCCCGAGAACATGAGGAGTTTC |
| GTR2_qR | TCAGGCAACGTTCTTGTCTTG |
| GTR3_qF | CAATCCGATAGACGCCTTGG |
| GTR3_qR | CAACCACATACCGGACATCG |
| UBQ2_qF | CCAAGATCCAGGACAAAGAAGGA |
| UBQ2_qR | TGGAGACGAGCATAACACTTGC |
